# Predicted Effector Gene Aggregation, Standards and Unified Schema (PEGASUS): A Community Framework for Effector Gene Reporting

**DOI:** 10.64898/2026.06.16.731894

**Authors:** Aoife McMahon, Yue Ji, Julie A. Jurgens, Maria C. Constanzo, Adam S. Butterworth, Matthew C. Pahl, Szymon Szyszkowski, Karl Helibron, Abdurrahman Shiyanbola, Yakov Tsepilov, Cassandra N. Spracklen, Jeremy A. Arbesfeld, Drew Hite, Alex Shilin, Elizabeth Lewis, PEG Working Group, Helen Parkinson, Noël P. Burtt, Laura W. Harris

## Abstract

Genome-wide association studies (GWAS) increasingly report predicted effector genes (PEGs) - genes hypothesised to mediate the biological effects of associated variants. These function as key outputs for advancing variant-to-function research, mechanistic understanding, and therapeutic discovery. However, the rapid growth of PEG lists has not been matched by standards for organising, annotating, and reporting these predictions. As shown by recent landscape analyses, PEG lists vary widely in methodology, evidence definition, nomenclature, provenance tracking, and data structure, limiting interoperability, benchmarking, reuse, and adherence to FAIR principles.

To address this gap, we convened an international multi-stakeholder community comprising method developers, data generators, resource maintainers, curators, funders, journal editors, and downstream users. Through a 2024 workshop and a 2025 working group series, we developed the Predicted Effector Gene Aggregation, Standards and Unified Schema (PEGASUS) framework. PEGASUS specifies (i) a metadata standard to capture provenance, trait and GWAS descriptors, evidence sources, and integration methods; (ii) a structured evidence matrix reporting all genes and all evidence underpinning prioritisation at each locus; and (iii) a concise PEG list that summarises author-prioritised genes linked transparently to underlying evidence. The framework balances transparency, machine readability, burden on submitters, and alignment with existing community standards.

PEGASUS provides the first community-developed schema for reporting predicted effector genes and their supporting evidence. Adoption of this framework by authors will improve the comparability, reproducibility, and reusability of PEG outputs across studies, facilitating more robust biological inference, enabling cross-resource comparison of gene-prioritisation methods to support community benchmarking, and integration into downstream resources and analytical pipelines. PEGASUS-compliant data can be shared via the PEG Data Registry platform (https://kpndataregistry.org/peg), promoting re-use and establishing the basis for future integration with publicly shared GWAS data.

## Introduction

Genome-wide association studies (GWAS) have transformed human genetics by identifying thousands of genomic loci associated with common diseases and traits. However, translating these statistical associations into biological insights remains a major challenge. Most GWAS loci encompass multiple genes and regulatory elements, and the variants driving the associations often have modest effects, act through non-coding mechanisms, and influence tissues or contexts that may be difficult to assay experimentally. As a result, determining the specific *effector gene(s)*—the gene (or genes) whose disruption or modulation mediates the trait-associated signal—is a critical but complex step in the variant-to-function (V2F) process.

In recent years, the genetics community has increasingly recognised the importance of reporting predicted effector genes (PEGs), also referred to as causal/candidate genes, as a core outcome of GWAS. Researchers commonly integrate many evidence types—e.g. genomic proximity, epigenomic annotations, expression and chromatin quantitative trait loci (QTLs), perturbational data, burden tests, literature evidence, and machine-learning-based prioritisation—to identify effector genes at associated loci. PEG lists have become widely used as starting points for functional follow-up studies, drug target discovery, mechanistic interpretations of GWAS results and training of gene-prioritisation models. However, as highlighted in recent landscape analyses (Costanzo et al, 2025), the rapid rise in PEG reporting has occurred without corresponding standards for describing, structuring, or sharing these predictions.

The absence of shared standards has led to substantial heterogeneity in how PEGs are generated and reported across studies. Evidence categories are defined inconsistently, methods for evidence integration vary widely, and the level of reporting detail ranges from full matrix-like summaries of all genes and evidence at each locus to minimal lists of top-ranked genes. PEG lists are frequently provided only as supplemental tables; information about evidence provenance, variant definitions, trait descriptors, or genome builds may be incomplete; and key metadata are often omitted or embedded in narrative text rather than structured fields. These inconsistencies hinder reproducibility, complicate benchmarking of gene-prioritisation methods, reduce interoperability with computational pipelines, and diminish adherence to FAIR (Findable, Accessible, Interoperable, and Reusable) principles (Wilkinson et al, 2016). They also limit the ability of downstream users to interpret and reuse PEG results. A recurring motivation for PEG standardisation is the increasing use of PEG outputs as inputs for computational and machine learning methods. Such approaches require structured, complete and consistently formatted data in order to aggregate results across studies, traits and methods and to train, validate and benchmark computational approaches. Standardised PEG outputs therefore provide an essential foundation for developing robust, AI-ready resources for variant to function research.

Recognising both the impact of PEGs and the challenges posed by their inconsistent reporting, we convened a broad community of stakeholders to explore the requirements for a shared standard. Here, we introduce the output of this working group: the Predicted Effector Gene Aggregation, Standards and Unified Schema framework (PEGASUS). PEGASUS provides a structured, community-developed standard for reporting predicted effector genes and the evidence supporting them. It defines:

(1) a PEG evidence matrix that systematically reports all genes evaluated at each locus and all supporting evidence, structured into evidence streams which are defined and described in the metadata; (2) a concise PEG list summarising an author’s prioritised genes, linked transparently to the underlying evidence; and (3) a metadata standard capturing evidence sources, integration approaches, provenance, trait descriptors and GWAS details. This structure can be seen as analogous to GWAS data, with (1) full genome-wide summary statistics and (2) the lead associations typically reported in the main tables of a paper after significance thresholding, LD-pruning and/or conditional analysis. Both datasets are supported by metadata including critical information required for interpretation and reuse

There is no agreed-upon single best way to make a PEG list, and the intention of PEGASUS is not to define one; the goal of this work is simply to foster transparency. By establishing reporting standards, PEGASUS addresses long-standing challenges in PEG interpretation and provides a foundation for reusability, reproducibility, benchmarking, and integration into downstream analytical and biological workflows. The framework represents the first community-driven effort to harmonise the reporting of predicted effector genes and is intended as a minimum data requirement to be applied in manuscript preparation as well as serve as a submission format. It is a living standard that will evolve alongside new data types, methods, and community needs.

## Methods

### Community-driven development process

The PEGASUS framework was created through a structured, community-led process coordinated by the Knowledge Portal Network (https://hugeamp.org/r/kpn_home) at the Broad Institute of MIT and Harvard and the NHGRI-EBI GWAS Catalog (www.ebi.ac.uk/gwas; Cerezo et al, 2025) at EMBL-EBI. The process involved an international team of researchers who generate PEG lists, users of PEG data, developers of gene prioritisation methods, curators of major genomic data resources, experts in metadata and data standards, and representatives from journals and funding agencies. This group was established to ensure that the standard would be practical for an array of contributors and useful for downstream reusers.

The development process combined a landscape analysis, a multi-site workshop, iterative working group meetings, and benchmarking exercises. Each component contributed distinct information that shaped the design of PEGASUS.

### Landscape analysis

Our team initially conducted a systematic review of PEG lists published across complex trait and disease GWAS to understand current practices (Costanzo et al, 2025). This assessment documented wide variation in evidence types, naming conventions, metadata completeness and reporting formats, ranging from simple ranked lists to extensive but inconsistently organised tables. The review also generated a set of high level recommendations that emphasised the need for clear terminology, structured evidence annotation, explicit provenance, standard identifiers, transparent prioritisation criteria and consistently formatted tables that report all evaluated genes and evidence. These recommendations served as an early foundation for later standardisation efforts. The insights from this analysis helped to identify challenges that any reporting standard would need to address and informed the agenda for the subsequent workshop.

### Workshop and working group activities

An international workshop held in September 2024 brought together contributors and users of PEG lists from academia, industry and major data resources. Presentations, surveys and breakout sessions were used to gather community perspectives on terminology, evidence requirements, metadata needs and barriers to reproducibility (Supplemental File 1). Participants identified common pain points, including unclear evidence definitions, inconsistent metadata, inadequate provenance tracking, and barriers to reusability, and expressed strong support for developing a flexible but structured community standard for PEG reporting.

As part of the workshop, an initial draft proposal for a PEG reporting standard (based on the findings of the landscape analysis) was presented as a starting point for discussion. This proposal outlined a minimal set of fields for reporting PEG lists and associated metadata and was explicitly intended to surface limitations and stimulate community feedback rather than to define a final specification. Feedback on this initial proposal informed subsequent design decisions, including the decision to support flexible integration strategies rather than imposing a single scoring framework.

A central conceptual insight emerged during these workshop discussions. Participants noted that PEG outputs naturally separate into two distinct products that serve different purposes: (1) a comprehensive evidence matrix containing all genes and all supporting evidence considered at each locus, and (2) a summarised PEG list reporting prioritised genes and intended for downstream biological use. This can be seen as analogous to the (1) full genome-wide summary statistics from a GWAS, and the (2) “top associations” that are typically reported in main tables of paper, both considered important outputs for different use cases. This distinction had not been formalised in earlier work, including the landscape review, and the workshop identified it as a critical structural element for any reporting standard.

Following the workshop, a standing working group met regularly throughout 2025 to formalise this distinction and to develop the technical components of the standard. The group refined metadata requirements, evidence category definitions, variant and locus representation, controlled vocabularies and file formats. Iterative drafts of the schema and templates were circulated and updated based on community feedback.

### Design principles, usability and implementation testing

Across all phases of development, the working group adhered to a set of guiding principles intended to balance flexibility with structure. These were (1) adherence to FAIR principles, (2) transparency in evidence provenance, (3) machine readability through standard structured formats, (4) interoperability with existing data resources and ontologies, (5) minimal and clear mandatory fields and (6) accommodation of multiple PEG prediction methods without imposing a single scoring framework. These principles shaped decisions about metadata structure, evidence category definitions, file formats and the distinction between the evidence matrix and summarised PEG list.

To assess whether the draft PEGASUS specification met these principles in practice, volunteers reformatted existing PEG lists from published studies into the proposed structure.

## Results

### Overview of the PEGASUS framework

The PEGASUS framework establishes a structured standard for reporting predicted effector genes and the evidence used to prioritise them. It is designed to support transparent and reproducible communication of gene prioritisation decisions while accommodating a wide range of analytical approaches. PEGASUS consists of three linked components: (1) a structured metadata record that captures the contextual information required to interpret PEG outputs, (2) a comprehensive evidence matrix that reports all genes considered at each locus and all associated evidence and (3) a summarised PEG list that presents the authors’ prioritised genes and links them directly to the underlying evidence (Figure 1; Supplemental File 1: Figure S1).

**Figure 1.**
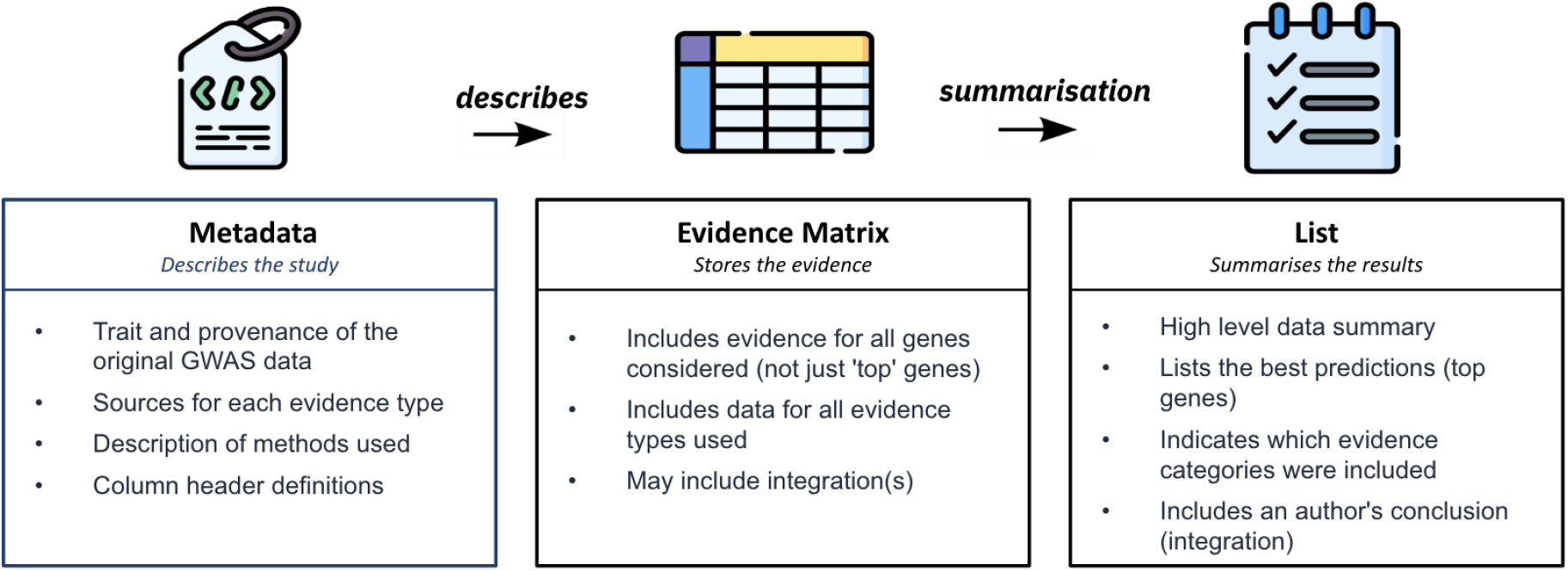
Overview of the PEGASUS framework. Schematic illustration of the three components of PEGASUS: metadata, evidence matrix and summarised PEG list, and how they relate to each other.

The framework operates at the level of a single trait or phenotype analysed through a GWAS or meta-analysis. Each GWAS dataset analysed for predicted effector genes produces one metadata record and one evidence matrix, along with a PEG list. This structure separates comprehensive evidence reporting from the summarised interpretation and supports many downstream use cases, including comparative method evaluation, knowledge base ingestion, machine learning applications, experimental design and drug discovery.

### Evidence matrix

The evidence matrix is the central data product of the PEGASUS framework. It provides a complete representation of the information used to evaluate candidate genes at each locus. The matrix is structured as a table in which each row corresponds to a specific variant-gene pair at a specific locus and each column corresponds to a defined evidence type, provenance field or integration metric (Figure 2A).

**Figure 2.**
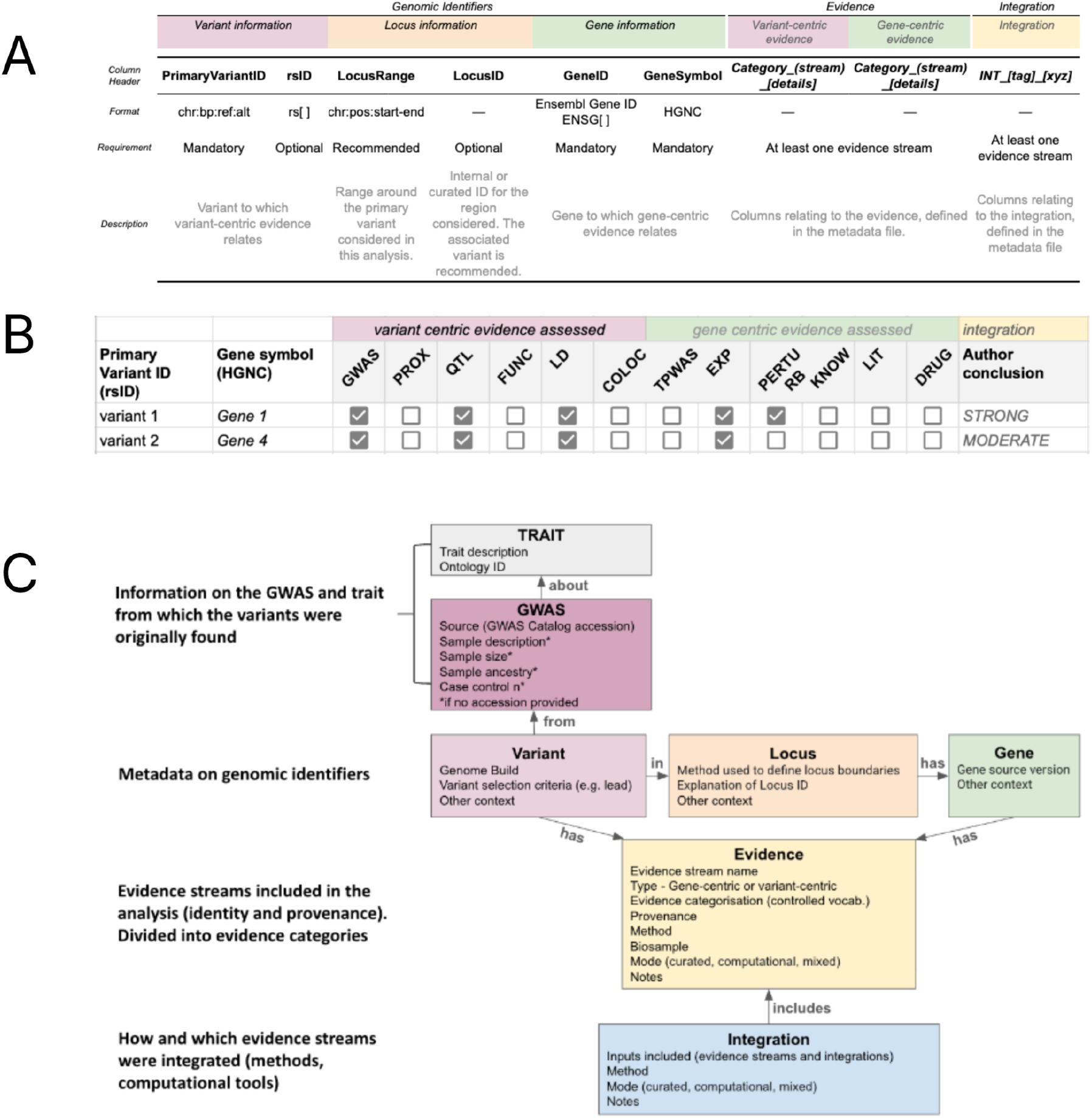
Representing predicted effector gene (PEG) list data under the PEGASUS standard. A. A representation of the evidence matrix that illustrates variant-centric evidence fields, gene-centric evidence fields, provenance columns and optional integration columns. The figure shows the standardised naming conventions and column groups used in PEGASUS. B. Each row in the PEG list represents a top gene per locus. Each column represents an evidence category, where the value indicates whether evidence was considered (tick = data present; blank = not assessed). The final column reflects the author’s integrated conclusion. C. A visual map of metadata fields grouped into major categories, including trait descriptors, GWAS provenance, locus definition, evidence definitions and integration description.

PEGASUS uses a controlled vocabulary to organise individual evidence streams into a defined set of evidence categories. To ensure consistent interpretation, each category’s abbreviation is used within column names and combined with other relevant metadata where needed. *Variant-centric evidence* includes information tied directly to specific GWAS variants, such as fine mapping probabilities, credible set membership or variant-level functional annotations. *Gene-centric evidence* includes attributes associated with genes, such as perturbation data, expression patterns, known disease associations or literature based annotations. This separation clarifies whether the evidence points primarily to the variant or the gene in each row.

The matrix allows the inclusion of integration fields when researchers choose to generate combined scores or prioritisation values. PEGASUS does not prescribe any specific method for combining evidence but requires that any integrated field be defined, accompanied by provenance information and a clear methodological description in the linked metadata. This flexibility ensures that authors can apply methods that best fit their analytical workflow while enabling downstream users to understand how prioritisation decisions were reached.

The evidence matrix supports both human and machine consumption. It can be used directly for downstream computational analyses, comparative benchmarking of gene prioritisation methods or automated ingestion into knowledge bases.

### Organisation of evidence categories

PEGASUS defines a structured vocabulary for evidence categories that distinguishes between two major evidence types: variant-centric evidence and gene-centric evidence. We grouped evidence types into these two broad categories to better represent the nature of the evidence and its relationship to gene prioritisation. This organisation is illustrated in Figure 4 and described in Supplemental File 1: Table S1. The use of a controlled vocabulary helps ensure consistent representation of evidence across studies and supports automated parsing of evidence matrices. It also improves clarity for downstream users by providing a shared framework for interpreting varied evidence types.

Variant-centric evidence includes, for example, methods that focus on statistical fine-mapping to identify likely causal variants. Once candidate variants are identified (method described in the metadata), additional annotations such as regulatory contacts, chromatin interaction data, QTLs, or known effects such as clinically relevant variants are used to infer gene effects at the locus level. This type of evidence directly connects genetic variants to their potential functional impact on genes. The full list of variant-centric evidence categories is shown in Supp Table 1a.

In contrast, gene-centric evidence considers the gene and its role. This category includes functional annotations such as gene expression data, perturbation experiments, gene set associations, gene burden analyses, and relevant literature knowledge. By focusing on the genes themselves, this approach integrates more holistic biological insights that may not be directly linked to specific variants but still contribute to understanding gene function and disease mechanisms. The full list of gene-centric evidence categories is shown in Supp Table 1b.

Importantly, the consistent categorisation of evidence enables integration and comparison across PEGASUS evidence matrices and derived PEG lists. By using shared evidence category definitions, PEGASUS supports aggregation of results across studies, methods and traits, and allows downstream users to interpret which types of evidence contributed to gene prioritisation in a comparable way. This is particularly important for applications that combine or contrast PEG outputs across multiple analyses.

We recognise that the list of sub-categories is not exhaustive and may change in future as methods evolve. The framework is therefore designed to remain flexible: categories define the overall analysis type (e.g., QTL analysis, which captures molecular relationships with genetic variation), while streams (sub-categories) capture finer-grained signal types (e.g., expression QTLs [eQTLs], and splicing QTLs [sQTLs]). This structure allows PEGASUS to maintain a stable high-level framework while accommodating expanding and increasingly granular evidence types without requiring redesign of the core system.

### PEG list

The PEG list provides a concise summary of genes prioritised by the authors. It is intended for users who require a high-level overview of candidate genes without exploring the complete evidence matrix. Authors must explicitly specify which integration represents their final conclusion in the evidence matrix, ensuring that the basis for gene selection is transparent and reproducible (Figure 2B). The PEG list can then be derived directly from the evidence matrix by selecting the locus-gene(s) relationships specified by the author_conclusion column.

In addition to gene identifiers, the PEG list provides a high-level summary of which evidence categories were considered by the authors when forming their conclusion. This summary is intended to give users a rapid overview of the types of evidence evaluated, rather than to indicate the direction, strength or consistency of individual evidence streams. Users requiring detailed or quantitative interpretation of the evidence are expected to refer to the underlying evidence matrix.

The PEG list does not replace the evidence matrix. Instead, it communicates the authors’ prioritisation in a compact and interpretable form while preserving a direct link to the full set of evaluated genes, evidence and integration logic. This separation supports different use cases and user groups, including experimental and translational users who may focus on the PEG list for hypothesis generation and computational users who may work directly with the evidence matrix for benchmarking, reanalysis or method development. Future iterations of PEGASUS may extend the PEG list to support richer representations of evidence and, where appropriate, more formalised scoring or ranking schemes, while maintaining compatibility with the underlying evidence matrix.

### Metadata standard

To ensure that PEGASUS data products can be interpreted and reused consistently, the framework also defines a structured metadata standard.

The metadata specification in PEGASUS ensures that essential contextual information is captured in a consistent and machine readable format. Metadata fields describe the trait analysed and specify both controlled vocabulary identifiers and a human readable trait description (Figure 2C). They also record the GWAS on which the PEG determination is based, including accession identifiers, sample characteristics, genome build and references to primary publications.

The metadata record documents the rules used to define loci, including the locus definition method, window size or fine mapping approach, and the criteria used to assign genes to loci. It also describes the evidence considered in the PEG analysis, including assignment to evidence categories (Figure 4, Supplemental File 1: Table S1), the sources of evidence, and the approaches used to combine or score evidence and relevant biosample details. Any integration method applied to generate prioritisation scores or rankings is fully described. Column header definitions are also provided. The metadata record therefore provides the provenance and interpretive detail required for reproducibility and interoperability.

PEGASUS metadata fields are divided into mandatory and recommended categories. Mandatory fields capture the information required to interpret any PEG output at a basic level (*e*.*g*., genome_build is essential since variant positions vary across different reference genome assembly versions); while recommended fields improve transparency and enable linkage with external resources and ontologies (*e*.*g*., variant_type indicates the variant selection criteria considered for gene prioritisation). To maximise interoperability, metadata are provided in a machine readable format such as YAML or JSON.

### Worked example with synthetic and real data

To illustrate the application of the PEGASUS framework, a synthetic dataset is provided in the PEGASUS documentation (https://gwas-catalog.github.io/PEGASUS/) and as Supplemental File 2. This dataset demonstrates how raw PEG-relevant information can be transformed into a complete metadata record, evidence matrix and PEG list. Figure 3 shows the progression from the initial synthetic data (panel A), to the metadata representation (panel B), and finally to the fully structured evidence matrix (panel C). The complete metadata file for the synthetic dataset is available in the Supplemental Material in YAML format, and the corresponding PEG list is shown in Figure 2B. This worked example allows users to understand the mechanics of the standard, observe how evidence fields map into the matrix structure and verify that their own outputs conform to the expected format.

**Figure 3.**
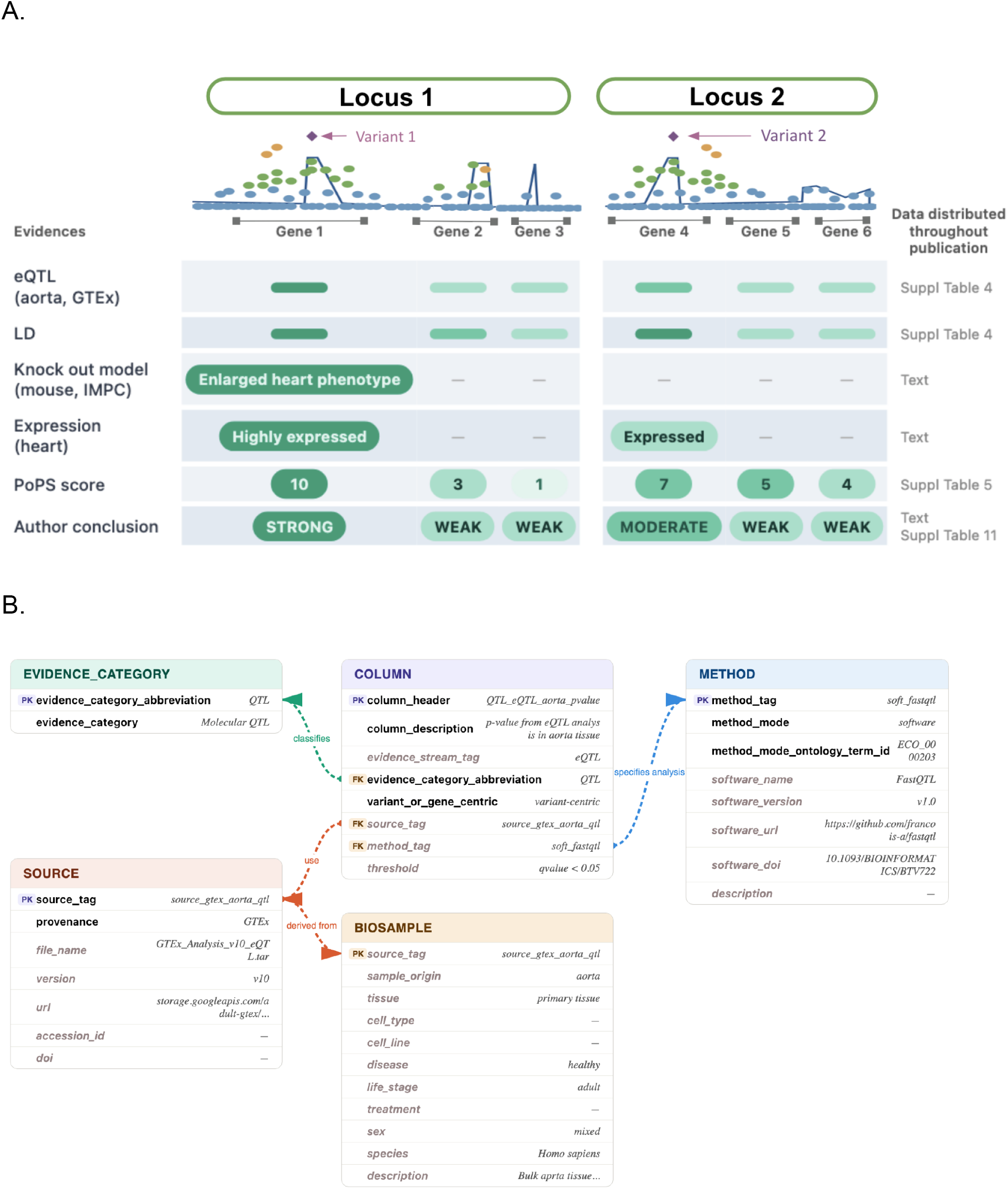
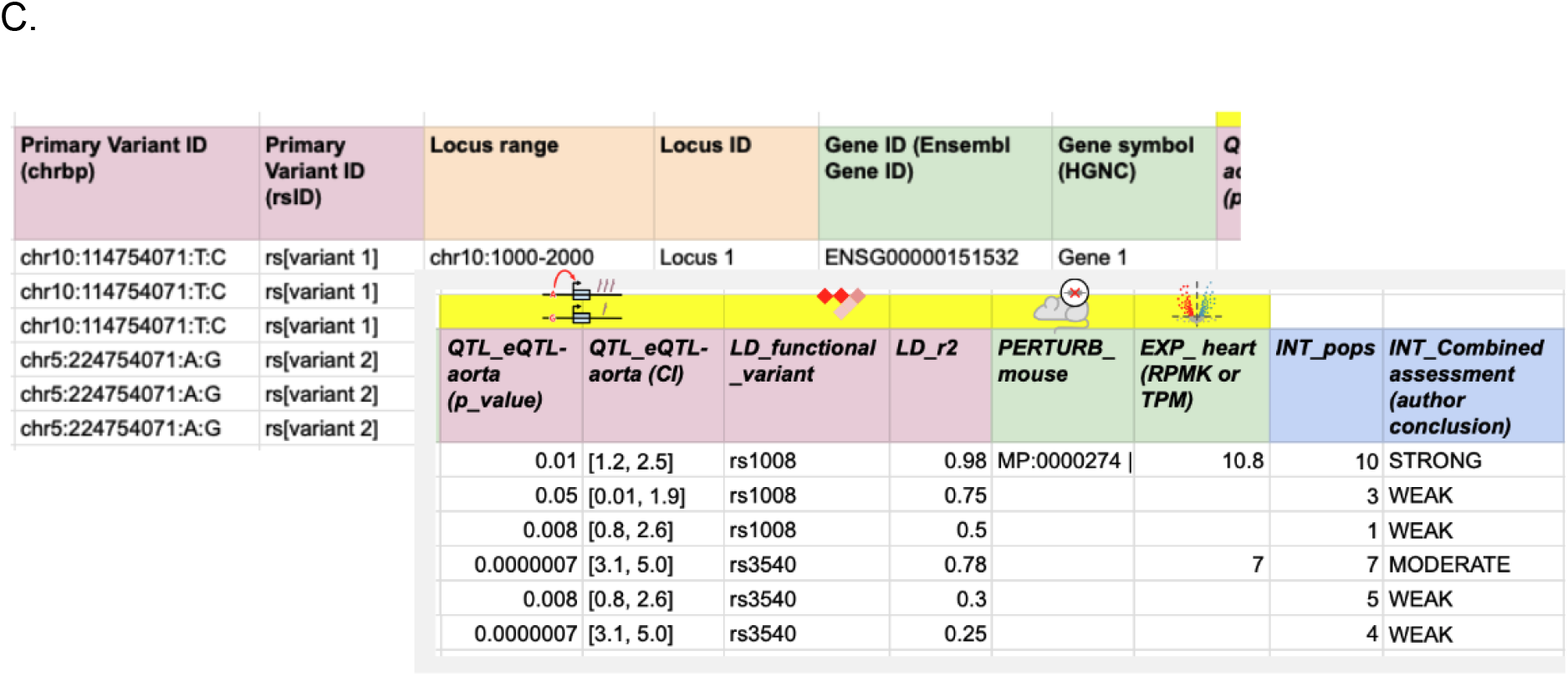
Worked example with synthetic data. A worked example demonstrating the PEGASUS framework. A. The initial synthetic dataset in its raw format. B. Entity–relationship diagram of the QTL evidence in the PEGASUS framework. The COLUMN entity defines measurable data features (e.g., p-values) linked to key metadata. Each column is classified by an EVIDENCE_CATEGORY, generated by a METHOD (e.g., FastQTL), and provided by a SOURCE (e.g., GTEx), which is further characterised by a BIOSAMPLE capturing biological context such as tissue, disease state, and species. Arrows denote relationships; Mandatory fields are shown in regular text; optional fields in lighter/italic text.;Primary key (PK) indicates the unique Identifier and foreign key (FK) provides the link between the tables. C. The PEGASUS evidence matrix derived from the synthetic dataset. Figure 2B shows the PEG list representation of the same synthetic dataset.

**Figure 4.**
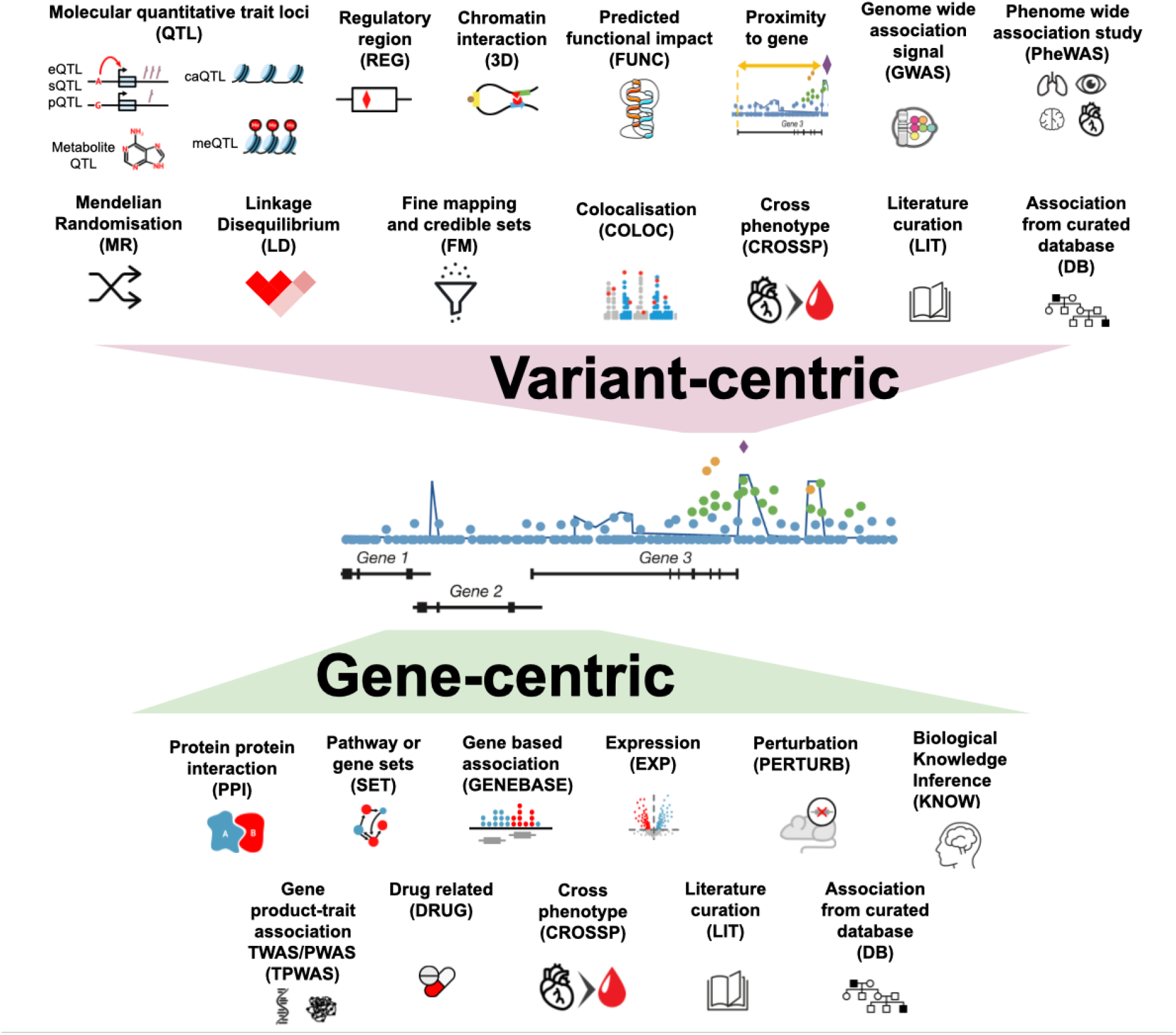
Evidence category organisation. Diagram showing the organisation of variant-centric and gene-centric evidence categories and the controlled vocabulary used to classify evidence types within the PEGASUS framework.

In addition to the synthetic dataset, PEGASUS has been applied to multiple published PEG outputs during working group benchmarking and community testing. As an illustrative example, a single locus from Aragam et al. (2022) was reformatted into PEGASUS-compliant metadata, evidence matrix and PEG list structures (Supplemental File 3). This example demonstrates how PEGASUS can represent existing gene prioritisation outputs by capturing reported evidence and integration results, without recomputing or reinterpreting the original analysis.

### Implementation outcomes and experience

Implementation exercises were conducted by members of the PEGASUS working group to assess the feasibility and usability of the framework across a range of PEG outputs. Participants applied the PEGASUS specification to previously published studies as well as to their own PEG analyses, allowing evaluation across different evidence types, traits and integration strategies.

Overall, participants reported that PEGASUS was sufficiently flexible to accommodate all tested datasets without requiring changes to the underlying analyses. The process of converting existing PEG outputs into PEGASUS-compliant files was generally completed within a reasonable timeframe, comparable to the effort required to submit data to established genomic data repositories. These findings indicate that the framework can be adopted in practice without imposing excessive burden on data generators. Feedback from these activities informed targeted refinements to the specification and documentation, further improving clarity and usability. Participants also noted that lightweight tooling, templates and validation tools to facilitate metadata entry would be valuable, particularly to support consistent population of metadata fields and to reduce manual effort during submission.

When applying PEGASUS to legacy outputs generated without standardisation in mind, some additional effort was required to interpret incomplete provenance, missing contextual information or bespoke evidence descriptions. These challenges largely reflected the curatorial nature of reformatting data produced by others, rather than limitations of the framework itself. In contrast, participants who tested PEGASUS using their own PEG outputs generally found the conversion process to be straightforward, reflecting access to full study context, provenance and intermediate evidence. In both cases, minor uncertainty when mapping authors’ evidence descriptions into structured fields highlighted areas where additional guidance or worked examples could further support consistent use.

Taken together, the exercises demonstrated that PEGASUS can be applied efficiently and consistently across heterogeneous PEG analyses. These results support the practical feasibility of PEGASUS as a standard reporting framework for predicted effector gene analyses, promoted by development of submission tools and infrastructure for data sharing.

### Data sharing

The PEGASUS framework, including examples of application, is available on GitHub (see Data availability section). We encourage authors and generators of PEG data to explore the framework and apply it in their published data. A template to support preparation of metadata is available as Supplemental File 4 and in the documentation.

PEGASUS-compliant data can be submitted to the PEG Data Registry platform (https://kpndataregistry.org/peg), and made available via the Predicted Effector Gene Knowledge Portal (www.pegkp.org).

## Discussion

### PEGASUS as a flexible and FAIR reporting framework

PEGASUS provides a community-developed framework for standardising the reporting of predicted effector genes and the evidence used to prioritise them. By defining a structured metadata record, a comprehensive evidence matrix and a summarised PEG list, PEGASUS addresses long-standing inconsistencies in how PEG outputs are described, formatted and shared across studies. The framework enables researchers to document both the underlying evidence and the interpretive decisions made during gene prioritisation, improving transparency, reproducibility and reuse.

A central strength of PEGASUS is its ability to accommodate varied gene prioritisation approaches while preserving consistent reporting. PEG studies vary widely in their use of GWAS designs, locus definition strategies, evidence types and integration methods. Rather than prescribing a single analytical workflow or scoring scheme, PEGASUS focuses on standardising how results and supporting information are reported. This approach allows methodological diversity to be preserved while enabling downstream users to understand how prioritisation decisions were reached and to compare outputs across studies, supporting the data to be reused in downstream applications.

Adopting an existing evidence ontology for the evidence categories defined in the PEGASUS schema facilitates FAIR data principles, particularly interoperability and reusability, because the ontology serves as an external resource providing standardised terms accompanying definitions that enable consistent interpretation. After reviewing the bioscientific data analysis ontology (EDAM) (Ison et al, 2013), the Evidence and Conclusion Ontology (ECO) (Nadendla et al, 2022), and the Ontology for Biomedical Investigations (OBI) (Bandrowski et al, 2016) as candidate ontologies, ECO emerged as the closest match to PEGASUS’s evidence categories. However, the mapping is not yet complete: several PEGASUS-specific evidence types lack direct ECO equivalents, and resolving these gaps will require further refinement of term definitions, which is a high priority for future work.

PEGASUS is explicitly aligned with FAIR data principles. The metadata schema improves findability and reusability by capturing stable identifiers, ontology terms and provenance information. The evidence matrix supports interoperability by using controlled vocabularies and machine readable structures that can be ingested by computational workflows and data resources. The separation between the evidence matrix and the summarised PEG list improves accessibility for different audiences, supporting both detailed computational reuse and high level human interpretation. Use cases include ingestion of data into downstream resources such as the GWAS Catalog and Predicted Effector Genes Knowledge Portal (PEGKP), enabling users to easily access PEGs defined for a GWAS locus. Currently, the GWAS Catalog includes mapped genes (*i*.*e*., the closest gene to a variant) only, whereas inclusion of PEGs would provide a richer annotation to inform downstream analysis. PEGKP has ingested a number of manually-curated PEG lists in order to provide comparative evidence to automated tools such as the Human Genetics Evidence (HuGE) Calculator, but these are limited in scope due to data missing at source (Costanzo et al, 2025). For PEG data to be efficiently ingested into resources, minimum FAIR reporting standards need to be applied, in order to ensure consistency and demonstrate provenance of the information. As a future direction, we will mint persistent identifiers and create landing pages for PEG data within the Knowledge Portal Network, enabling efficient access and reuse by both humans and agentic systems (Supplemental File 1: Figure S2). Our overall implementation decisions, prioritizing adherence to FAIR principles, will ensure that PEGASUS is broadly accessible to researchers across a wide range of scientific areas.

### Towards ‘gold standard’ gene sets

Widely accepted gold standard sets of effector genes–*i*.*e*., those which have been consistently replicated and manually verified within a given trait–do not yet exist, but PEGASUS is designed to support their development. Even the current PEGASUS list representation enables pragmatic approaches to constructing reference gene sets. When multiple PEGASUS-compliant lists are generated for the same trait using independent methods or datasets, simple overlap-based aggregation can be used to identify genes that are consistently prioritised. Such approaches provide an initial and transparent means of defining provisional reference sets that can be used for benchmarking, method evaluation and as training sets for machine learning methods (for example, the Open Targets L2G pipeline (Mountjoy et al, 2021; Tsepilov et al, 2026)

In the longer term, PEGASUS also provides a foundation for more refined approaches to gold standard generation. As PEGASUS lists accumulate across traits, it becomes feasible to explore more systematic scoring or weighting strategies that combine evidence across studies. These efforts could draw on lessons from other structured scoring frameworks in human genomics (Strande 2017), which pair explicit evidence criteria with expert community review. Complementary community curation by trait specific domain experts could enable iterative refinement of reference gene sets as biological knowledge evolves. Together, these approaches position PEGASUS not only as a reporting framework but as infrastructure that can support the development of increasingly robust, community endorsed gold standard resources and can provide the basis for identification of consensus biological mechanisms and novel insights.

### Future directions and community stewardship

We encourage authors and generators of PEG data to explore the framework, apply it in their published data and share via the PEG Data Registry platform (https://kpndataregistry.org/peg) to make available through the Predicted Effector Gene Knowledge Portal (www.pegkp.org). In the future, we aim to add reciprocal links between this database and the GWAS Catalog, enabling users to easily find PEG lists derived from GWAS datasets, and to access the underlying GWAS data for a given PEG list. Submission of data will promote their re-use in downstream applications such as target identification, facilitate replication and help to establish best practices in analytical methods.

PEGASUS is intended as a living framework that can evolve alongside advances in GWAS analysis, functional genomics and gene prioritisation methods. Future development may include creation of tools to support validation and submission, improved guidance and automation for metadata generation and closer alignment with emerging ontologies and data standards. As community practice matures, future iterations of the PEGASUS list may explore richer representations of evidence, such as indicating the relative strength or consistency of supporting evidence or explicitly capturing negative or conflicting evidence. Additional extensions could include author-defined thresholding strategies for PEG list inclusion, allowing genes to be reported based on explicit criteria rather than selecting a single top ranked gene per locus. Ongoing community engagement will be essential to ensure that PEGASUS remains responsive to new use cases and methodological developments, and we welcome feedback.

By establishing a common structure for reporting predicted effector genes, PEGASUS provides a foundation for improved transparency, reproducibility and reuse across variant-to-function research. Its adoption will support more robust comparison of gene prioritisation methods, clearer documentation of analytical choices and should support more effective translation of GWAS discoveries into biological insight.

## Acknowledgements

This work was supported in part by NIH/NHGRI U24HG011453 (N.B), NIH/NIDDK UM1DK105554 (N.B), NIH/NHGRI U24HG012542, European Bioinformatics Institute (EMBL-EBI) Core Funds, Open Targets, and by the Wellcome Sanger Institute core grant from Wellcome (220540/Z/20/A). The Cardiovascular Epidemiology Unit at the University of Cambridge has been supported by core funding from the British Heart Foundation (BHF) (RG/18/13/339463946: RG/F/23/110103), National Institute for Health and Care Research (NIHR) Cambridge Biomedical Research Centre (BRC-1215–20014: NIHR203312) [*], BHF Chair Award (CH/12/2/29428), Cambridge BHF Centre of Research Excellence (RE/24/130011, RE/18/1/34212) and by Health Data Research UK, which is funded by the UK Medical Research Council, Engineering and Physical Sciences Research Council, Economic and Social Research Council, Department of Health and Social Care (England), Chief Scientist Office Scottish Government Health and Social Care Directorate, Health and Social Care Research and Development Division (Welsh Government), Public Health Agency (Northern Ireland), British Heart Foundation and the Wellcome Trust.

*The views expressed are those of the authors and not necessarily those of the NIHR or the Department of Health and Social Care.

## Author contributions

AM, MCC, YT, CNS, HP, the PEG Working Group, NPB, and LWH conceptualized the study. AM, YJ, JAJ, MCC, CNS, NPB, and LWH developed methodology. AM, YJ, JAJ, MCC, EL and the PEG Working Group conducted investigations. AM, YJ, JAJ, and LWH wrote the original draft of the manuscript. AM, YJ, MCC, JAA, and LWH created visualizations. YJ, DH, AS and EL wrote software. YJ performed formal analysis. YJ and JAJ performed project administration. MCC performed data curation. ASB, MP, SS, KH, AS, and YT conducted data validation. ASB, MP, SS, KH, AS, YT, JAA, HP, and NPB reviewed and edited the manuscript draft. HP and NPB acquired funding. HP, NPB, and LWH performed project supervision.

## Data availability

The documentation, schemas, synthetic examples and benchmarking examples generated during this study are available at https://gwas-catalog.github.io/PEGASUS/. The workshop materials that helped inform the findings of this study are available at https://kp4cd.org/2024_PEG_workshop.

## Description of Supplemental files

1. **Supplemental File 1**: Contains the following supplemental items:
  - Summary of the September 2024 Community Workshop on Predicted Effector Gene Reporting, PEG working group membership.
  - Figure S1: Analogy between PEGASUS and commonly reported components of GWAS datasets.
  - Table S1: Evidence categories & definitions supported in the PEGASUS framework.
2. **Supplemental File 2: Synthetic_data_in_PEGASUS_schema.zip:** Synthetic example data presented in PEGASUS format, comprising matrix, metadata & list
3. **Supplemental File 3: Example_data_in_PEGASUS_Aragam.zip**: Real example data (Aragam et al, 2022) presented in the PEGASUS format.
4. **Supplemental File 4: Metadata_peg_template.xlsx**: Template to support preparation of metadata. Completed templates can be submitted to the PEG Data Registry platform (https://kpndataregistry.org/peg) and together with evidence matrices.

