## Supplemental Figure S1 for "Predicted Effector Gene Aggregation, Standards and Unified Schema (PEGASUS): A Community Framework for Effector Gene Reporting"

**Figure S1. Analogy between PEGASUS and commonly reported components of GWAS datasets**

| Function | PEGASUS component | GWAS |
| --- | --- | --- |
| - computational reuse - secondary analysis - detailed comparison of results between studies | PEG evidence matrix | Full genomewide summary statistics  (typically shared as via the GWAS Catalog or other repository, or as a supplementary file) |
| - communicating and interpreting the primary findings | PEG list | lead associations  (typically reported in the main tables of a paper after significance thresholding, LD-pruning and/or conditional analysis). |
| - critical supporting information required for interpretation and reuse | Metadata | |
