## Supplemental File 1 for "Predicted Effector Gene Aggregation, Standards and Unified Schema (PEGASUS): A Community Framework for Effector Gene Reporting"

### Supplemental File 1 Table of Contents

1. PEG Working Group Membership
2. Supplemental Note: Summary of the September 2024 Community Workshop on Predicted Effector Gene Reporting
3. Supplemental Note: Summary of the September 2024 Community Workshop on Predicted Effector Gene Reporting
4. Supplemental Figure S1. Analogy between PEGASUS and commonly reported components of GWAS datasets
5. Supplemental Figure S2. Workflow demonstrating PEGASUS accessibility to both humans and agentic AI systems
6. Supplemental Table S1. Evidence categories & definitions supported in the PEGASUS framework

#### PEG Working Group Membership

Beena Akolkar, Zhanna Balkhiyarova, Oleg Borisov, MacKenzie Brandes, Marcos Casado Barbero, Nathalie Chami, Daniel Considine, Ayse Demirkan, Eric Fauman, Xiangyu Ge, Quy Hoang, Emrah Kacar, Kanika Kanchan, Lillya Kopanitsa, Elizabeth Lewis, Yong Li, Ellen McDonagh, Florence Nakabiri, Nina Oparina, Santhi Ramachandran, Wafaa M. Rashed, Gabi Rinck, Oliver Ruebenacker, Elliot Sollis, Lorraine Southam, Sylvanus Toikumo, Kyle Vogan, Benjamin Wingfield, Juliana Xavier de Miranda Cerqueira, Sean Yao, Norann Zaghloul, Chi Zhang.

#### Supplemental Note: Summary of the September 2024 Community Workshop on Predicted Effector Gene Reporting

**Overview**

In September 2024, the Knowledge Portal Network and the NHGRI-EBI GWAS Catalog jointly convened a hybrid community workshop on standards and infrastructure for predicted effector gene (PEG) lists. The workshop brought together approximately 80 participants from academia, industry, data resources, method development, journal editorial teams and funding agencies. The primary aims were to assess current practices in PEG generation and reporting, identify barriers to reuse and interoperability and explore the feasibility of developing a community standard.

The workshop was held over two half days with in-person meetings at the Broad Institute and EMBL-EBI, alongside remote participation. Live note taking, pre- and post-workshop surveys and breakout discussions were used to capture community perspectives.

Workshop materials, including slides, meeting notes and recordings are available at <https://kp4cd.org/2024_PEG_workshop> and the PEGASUS Documentation in the Community section <https://ebispot.github.io/PEGASUS/docs/category/community>

**Workshop structure and scope**

Day one focused on the landscape of effector gene prediction, with presentations highlighting the diversity of gene prioritisation approaches and the challenges of interpreting GWAS signals across traits. Speakers from industry and large-scale resources described how PEG outputs are generated, curated and used in downstream applications such as drug target prioritisation and model training. Case studies illustrated the variability in evidence types, scoring strategies and reporting formats across published studies.

Day two focused on standardisation and reuse. Presentations covered prior experiences developing community standards, including GWAS summary statistics standards, and discussed the requirements for PEG lists to be reusable as inputs for computational methods, benchmarking efforts and knowledge graph applications. Results from a community survey were presented, followed by discussion of an initial “strawman” proposal for PEG reporting standards.

**Landscape challenges and motivations for standardisation**

Across sessions, participants consistently noted that PEG outputs are highly heterogeneous. Differences include how loci are defined, which genes are considered, which evidence types are included and how evidence is combined into prioritisation decisions. PEG outputs range from ranked gene lists to extensive supplementary tables, figures or images, often without clear provenance or structured metadata.

Participants agreed that this heterogeneity makes PEG outputs difficult to compare across studies, challenging to integrate into resources and unsuitable for many computational applications. The lack of consistent identifiers, evidence definitions and metadata was identified as a major barrier to FAIR reuse.

A recurring motivation for standardisation was the increasing use of PEG outputs as inputs for computational and machine learning methods. Participants highlighted that such methods require structured, complete and consistently formatted data in order to aggregate results across studies, traits and methods. Inconsistent reporting makes it difficult to determine which genes were evaluated at a locus, which evidence was considered and how prioritisation decisions were derived, limiting the ability to train, validate and benchmark computational approaches, where the inclusion of negative and uncertain examples is also informative.

Participants also discussed the absence of widely accepted gold standard or truth sets of effector genes, noting that computational and AI approaches depend on large, consistently described datasets to learn meaningful biological patterns. Standardised reporting was therefore seen as a prerequisite for developing robust, AI ready resources that can support deeper biological insight in variant to function research.

**Community survey insights**

Results from the pre-workshop community survey indicated broad agreement on several points. Respondents preferred the term “predicted effector gene” over “causal gene” when describing GWAS-based gene prioritisation outputs. Most respondents wanted access to evidence for all genes considered at a locus, not only the top prioritised gene. There was strong interest in being able to compare PEG outputs across studies and traits and in interacting with PEG data rather than consuming static tables.

Survey responses also highlighted the importance of transparency in how PEGs are generated, including explicit documentation of evidence sources, integration methods and assumptions.

**Initial proposal for a PEG reporting standard**

As part of the workshop, the organisers presented an initial proposal for a PEG reporting standard. This proposal was explicitly framed as a strawman standard, defined as a deliberately simple draft intended to stimulate discussion, identify shortcomings and encourage the development of improved alternatives, rather than as a final or prescriptive solution.

The strawman proposal outlined a minimal set of elements intended to support interoperability and reuse, including basic metadata to capture trait and GWAS provenance, a structured representation of evidence. It was intentionally incomplete and avoided prescribing specific evidence types, scoring systems or integration methods. Although the proposal was not developed in depth, it provided a concrete reference point that helped participants articulate concerns about overly restrictive standards, emphasise transparency over uniformity and clarify the need to support both comprehensive evidence reporting and concise summarised outputs. Feedback on the strawman helped refine the scope and direction of subsequent standard development.

**Key conceptual outcomes**

A central conceptual outcome of the workshop emerged during open discussion and breakout sessions. Participants recognised that PEG outputs naturally separate into two complementary products that serve different audiences and use cases:

1. A comprehensive evidence table or matrix that reports all genes evaluated at each locus together with all supporting evidence.
2. A summarised PEG list that reflects the authors’ prioritised genes and interpretive conclusions.

Participants agreed that these products should be treated as distinct but linked outputs and that standardisation efforts should explicitly support both. This distinction was not formalised in prior PEG landscape analyses and was identified by the workshop as a foundational design principle for any reporting framework.

**Breakout discussions and priorities**

Breakout groups discussed practical requirements for a PEG reporting standard. Common themes included:

- prioritising transparency over enforcing a universal scoring system,
- minimising mandatory fields while ensuring interpretability,
- supporting both human-readable summaries and machine-readable evidence matrices,
- clearly distinguishing newly generated experimental evidence from reused public data,
- enabling versioning and updates without imposing excessive burden,
- and providing guidance, examples and tooling to support adoption.

Participants also emphasised the importance of incentives for standard adoption, including alignment with journal requirements and deposition in trusted repositories.

**Relationship to existing standards**

Presentations on GWAS summary statistics standards and ClinGen frameworks highlighted the value of community engagement, iterative refinement and alignment with FAIR principles. These examples informed discussion about governance, versioning and long-term maintenance of PEG standards. Participants agreed that PEG lists are more complex than GWAS summary statistics but that similar community-driven processes could be effective.

**Outcome and next steps**

The workshop established strong community support for developing a PEG reporting standard and identified core requirements and design principles. It also led directly to the formation of a community working group, convened after the workshop, to translate these insights into a concrete framework. The PEGASUS framework described in the main manuscript is a direct outcome of this process.

#### **Supplemental Figure S1. Analogy between PEGASUS and commonly reported components of GWAS datasets**

| Function | PEGASUS component | GWAS |
| --- | --- | --- |
| - computational reuse - secondary analysis - detailed comparison of results between studies | PEG evidence matrix | Full genomewide summary statistics  (typically shared as via the GWAS Catalog or other repository, or as a supplementary file) |
| - communicating and interpreting the primary findings | PEG list | lead associations  (typically reported in the main tables of a paper after significance thresholding, LD-pruning and/or conditional analysis). |
| - critical supporting information required for interpretation and reuse | Metadata | |

#### **Supplemental Figure S2. Workflow demonstrating PEGASUS accessibility to both humans and agentic AI systems**

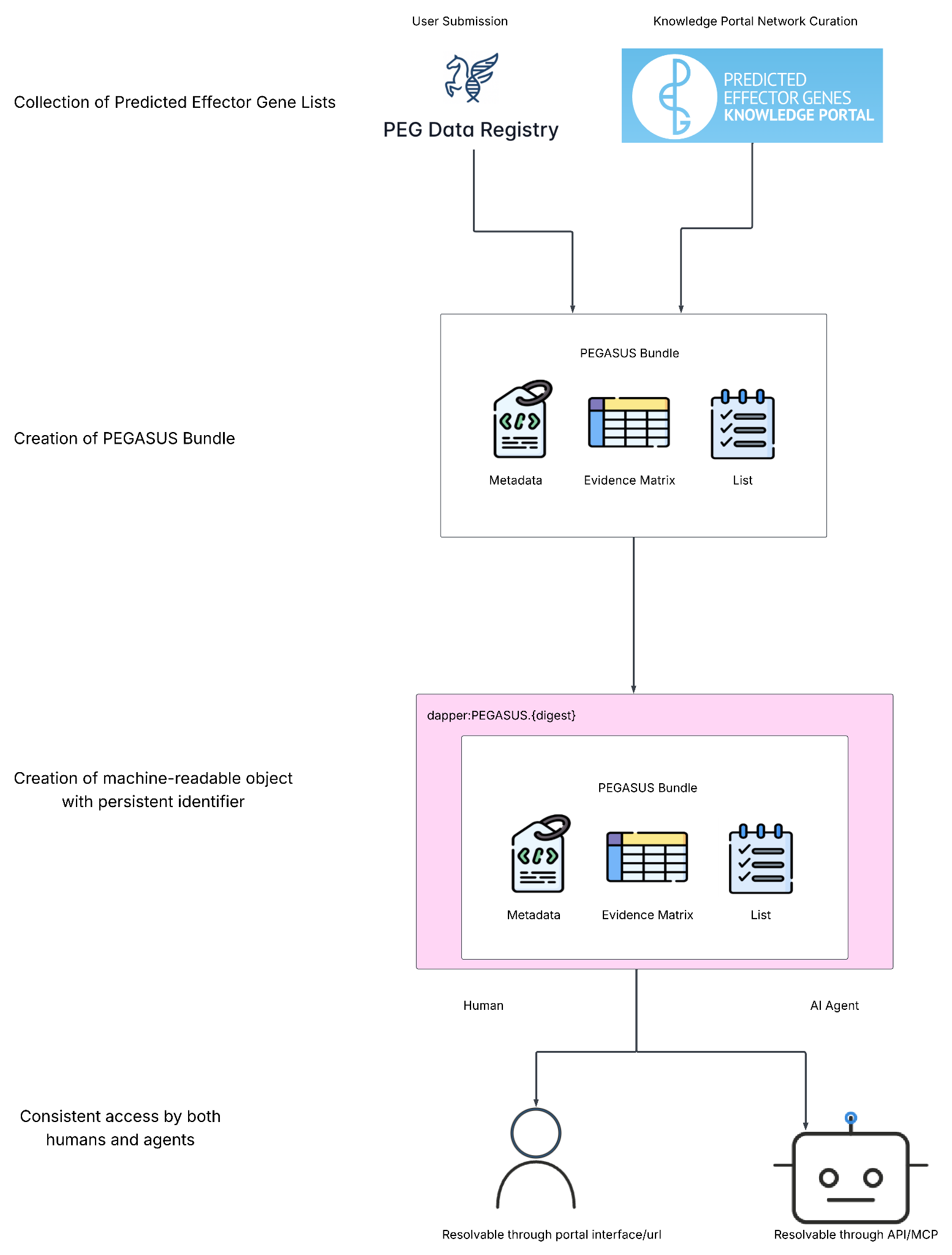

#### **Supplemental Table S1. Evidence categories & definitions supported in the PEGASUS framework**

| Supplemental Table S1a. Variant-centric evidence. | | |
| --- | --- | --- |
| Evidence categories | Abbreviation | Explanation |
| Linkage disequilibrium | LD | Assessment of whether a variant is correlated with another variant of interest and may act as a proxy. |
| Finemapping and credible sets | FM | Finemapping results - probability of variant being causal within a credible set, using Bayesian or probabilistic models |
| Colocalisation | COLOC | Variant affects two traits (typically a complex trait and a molecular phenotype) at the same locus. |
| Molecular QTL | QTL | Variant affects a molecular phenotype, e.g. gene expression (eQTL), splicing (sQTL), or protein expression (pQTL). |
| Mendelian Randomisation (MR) | MR | Uses genetic variants as proxies for exposures to test their causal effect on outcomes. |
| Regulatory region | REG | Variant lies in open chromatin or enhancer/promoter elements in relevant tissue (e.g. ATAC-seq, DNase-seq, or histone mark data) |
| Chromatin interaction | CHROMATIN | Variant lies in a region physically interacting with a gene promoter via 3D chromatin architecture (e.g. Hi-C, Capture-C data). |
| Predicted functional impact | FUNC | Variant predicted to disrupt gene/protein function or regulatory motifs, e.g. via SIFT, PolyPhen, CADD. |
| Proximity to gene (distance) | PROX | Assessment of whether variant is within or near gene boundaries. |
| Genome-wide association (GWAS) signal | GWAS | P-value from source GWAS for association of variant with trait specified in metadata file |
| PheWAS (Phenome-Wide Association Study) | PHEWAS | Variant is associated with multiple traits, suggesting pleiotropic effects |
| Cross-phenotype* | CROSSP | Gene or variant already established in a related phenotype (biologically similar). |
| Literature curation* | LIT | Human-curated gene or variant–disease links from literature. |
| Association from curated database* | DB | Variant or Gene is curated as causal or related to the phenotype from existing databases, like ClinVar, ClinGen, OMIM, etc. |

Evidence categories marked with * can also serve as gene-centric evidence

| Supplemental Table S1b. Gene-centric evidence | | |
| --- | --- | --- |
| Evidence categories | Abbreviation | Explanation |
| Protein–protein interaction | PPI | Gene’s protein interacts with other disease-relevant proteins. |
| Pathway or gene sets | SET | Gene is part of a known pathway or complex relevant to the phenotype, e.g. results of enrichment analyses using Reactome or KEGG. |
| Gene-based association | GENEBASE | Aggregated analysis of association of variants in gene with trait (e.g. SKAT, MAGMA, burden tests) . |
| Expression | EXP | Gene is differentially expressed in relevant tissue or disease e.g. the gene is more highly expressed in phenotype-related tissues compared to others. |
| Perturbation | PERTURB | Gene perturbation causes phenotype-relevant effects in lab or model organisms (knock out animal/cell line, human organoid). |
| Biological Knowledge Inference | KNOW | Gene–phenotype relationships can be inferred based on known biology, without providing specific references or direct experimental evidence linking the specific gene to the phenotype. |
| Genetically predicted trait association (TWAS/PWAS) | TPWAS | Evidence from transcriptome- or proteome-wide association studies showing that gene’s genetically predicted expression or protein level is associated with phenotype, |
| Drug related | DRUG | Evidence from drugs’ mechanism of action, e.g. gene encodes a known drug target or interacts with targets of drugs used to treat the phenotype, supporting therapeutic relevance. |
| Cross-phenotype* | CROSSP | Gene or variant already established in a related phenotype (biologically similar). |
| Literature curation* | LIT | Human-curated gene or variant–disease links from literature. |
| Association from curated database* | DB | Variant or Gene is curated as causal or related to the phenotype from existing databases, like ClinVar, ClinGen, OMIM, etc. |

Evidence categories marked with * can also serve as variant-centric evidence.
