## Supplemental File 3 for "Predicted Effector Gene Aggregation, Standards and Unified Schema (PEGASUS): A Community Framework for Effector Gene Reporting": metadata_Aragam_PEGSt000007.yml

```
DatasetDescription:
- trait_description: coronary artery disease
  trait_ontology_id: EFO_0001645
  peg_source: PMID:36474045
  gwas_source: GCST90132315
GenomicIdentifier:
- variant_type: significant
  variant_information: 279 significant associations from GWAS meta-analysis
  genome_build: GRCh37
  info: All protein-coding genes within 500 kb of all 279 genome-wide association
Evidence:
- column_header: GWAS_beta
  column_description: The effect size estimate from GWAS; indicates the direction
    and magnitude of the allele’s effect on the trait.
  evidence_category: Genome-wide association (GWAS) signal
  evidence_category_abbreviation: GWAS
  variant_or_gene_centric: variant-centric
- column_header: GWAS_pvalue
  column_description: The p-value of association between the variant and the trait
    from GWAS.
  evidence_category: Genome-wide association (GWAS) signal
  evidence_category_abbreviation: GWAS
  variant_or_gene_centric: variant-centric
- column_header: PROX
  column_description: If the gene is the nearest gene to the CAD sentinel variant.
  evidence_category: Proximity to gene (distance)
  evidence_category_abbreviation: PROX
  variant_or_gene_centric: variant-centric
  note: "Summary of the tissue-specific enrichment and functionally-informed fine-mapped\
    \ for the 127 regions enriched in at least one of the ten enriched CAD-relevant\
    \ tissues. Enrichments showing a >3SD increment in Bayes Factor (BF) are highlighted\
    \ in red. \nNULLBF = Bayes Factor under the null model; BESTBFJUMP = maximum increment\
    \ (i.e. enrichment) observed across the ten tissues; PPA = posterior probability\
    \ of association; HUVEC = human umbilical vein endothelial cells; HSMM = human\
    \ skeletal muscle myoblasts."
- column_header: FUNC_VEP_consequence
  column_description: Functional consequence of the variant predicted by Ensembl VEP.
    This column includes only protein altering consequence.
  evidence_category: Predicted functional impact
  evidence_category_abbreviation: FUNC
  variant_or_gene_centric: variant-centric
  method_tag: soft_vep
  threshold: protein-altering
  method:
    method_tag: soft_vep
    method_mode: software
    method_mode_ontology_term_id: ECO_0000203
    software_name: Ensembl VEP
    software_version: v96.0
    software_url: https://www.ensembl.org/info/docs/tools/vep/index.html
- column_header: QTL_eQTL_summary
  column_description: Binary indicator of whether there is evidence of a significant
    eQTL association in any evaluated GTEx or STARNET tissue. A value of 1 indicates
    that at least one GTEx or STARNET tissue showed a significant eQTL association
    for the locus/gene; a value of 0 indicates that no significant eQTL association
    was identified in the evaluated tissues.
  evidence_stream_tag: eQTL
  evidence_category: Molecular QTL
  evidence_category_abbreviation: QTL
  variant_or_gene_centric: variant-centric
- column_header: QTL_eQTL_gtex_Adipose_Subcutaneous_pvalue
  column_description: Nominal P-value for the association between genotype and gene
    expression in GTEx subcutaneous adipose tissue.
  evidence_stream_tag: eQTL
  evidence_category: Molecular QTL
  evidence_category_abbreviation: QTL
  variant_or_gene_centric: variant-centric
  source_tag: source_eqtl_gtex_AdiposeSubcutaneous
  method_tag: soft_eqtl
  threshold: FDR<5%
  note: 'eQTL look-ups from CAD-relevant tissues (adipose [subcutaneous, visceral
    omentum], adrenal gland, artery [aorta, coronary, tibial], liver and whole blood)
    in GTEx v7. Associations are only shown where the eQTL reached a 5% false-discovery
    rate threshold and the lead eQTL variant was in high LD (r2≥0.8) with the CAD
    sentinel variant.  Nominal P values were estimated using a two-tailed t-test.

    eQTL = expression quantitative trait locus.'
  source:
    source_tag: source_eqtl_gtex_AdiposeSubcutaneous
    provenance: GTEx
    version: v7
    url: https://www.gtexportal.org/home/datasets
    sample_origin: primary-tissue
    tissue: Adipose Subcutaneous
    cell_type: mixed
    life_stage: adult
    sex: mixed
    species: human
  method:
    method_tag: soft_eqtl
    method_mode: software
    method_mode_ontology_term_id: ECO_0000203
    method_description: 'We cross-referenced the sentinel CAD variants and their proxies
      (r2≥0.8) with STARNET eQTLs

      reaching a 5% FDR for all tissues. To ensure the CAD association and eQTL are
      likely to be

      driven by the same causal variant, we retained only those eQTLs where the CAD-associated

      variant and the lead eQTL variant had r2≥0.8 among European ancestry participants
      from UK

      Biobank. '
- column_header: QTL_eQTL_gtex_Adipose_Visceral_pvalue
  column_description: Nominal P-value for the association between genotype and gene
    expression in GTEx visceral adipose tissue.
  evidence_stream_tag: eQTL
  evidence_category: Molecular QTL
  evidence_category_abbreviation: QTL
  variant_or_gene_centric: variant-centric
  source_tag: source_eqtl_gtex_AdiposeVisceral
  method_tag: soft_eqtl
  threshold: FDR<5%
  source:
    source_tag: source_eqtl_gtex_AdiposeVisceral
    provenance: GTEx
    version: v7
    url: https://www.gtexportal.org/home/datasets
    sample_origin: primary-tissue
    tissue: Adipose Visceral
    cell_type: mixed
    life_stage: adult
    sex: mixed
    species: human
  method:
    method_tag: soft_eqtl
    method_mode: software
    method_mode_ontology_term_id: ECO_0000203
    method_description: 'We cross-referenced the sentinel CAD variants and their proxies
      (r2≥0.8) with STARNET eQTLs

      reaching a 5% FDR for all tissues. To ensure the CAD association and eQTL are
      likely to be

      driven by the same causal variant, we retained only those eQTLs where the CAD-associated

      variant and the lead eQTL variant had r2≥0.8 among European ancestry participants
      from UK

      Biobank. '
- column_header: QTL_eQTL_gtex_Adipose_Visceral_Omentum_pvalue
  column_description: Nominal P-value for the association between genotype and gene
    expression in GTEx visceral omentum adipose tissue.
  evidence_stream_tag: eQTL
  evidence_category: Molecular QTL
  evidence_category_abbreviation: QTL
  variant_or_gene_centric: variant-centric
  source_tag: source_eqtl_gtex_AdiposeVisceralOmentum
  method_tag: soft_eqtl
  threshold: FDR<5%
  source:
    source_tag: source_eqtl_gtex_AdiposeVisceralOmentum
    provenance: GTEx
    version: v7
    url: https://www.gtexportal.org/home/datasets
    sample_origin: primary-tissue
    tissue: Adipose Visceral Omentum
    cell_type: mixed
    life_stage: adult
    sex: mixed
    species: human
  method:
    method_tag: soft_eqtl
    method_mode: software
    method_mode_ontology_term_id: ECO_0000203
    method_description: 'We cross-referenced the sentinel CAD variants and their proxies
      (r2≥0.8) with STARNET eQTLs

      reaching a 5% FDR for all tissues. To ensure the CAD association and eQTL are
      likely to be

      driven by the same causal variant, we retained only those eQTLs where the CAD-associated

      variant and the lead eQTL variant had r2≥0.8 among European ancestry participants
      from UK

      Biobank. '
- column_header: QTL_eQTL_gtex_Adrenal_Gland_pvalue
  column_description: Nominal P-value for the association between genotype and gene
    expression in GTEx adrenal gland tissue.
  evidence_stream_tag: eQTL
  evidence_category: Molecular QTL
  evidence_category_abbreviation: QTL
  variant_or_gene_centric: variant-centric
  source_tag: source_eqtl_gtex_AdrenalGland
  method_tag: soft_eqtl
  threshold: FDR<5%
  source:
    source_tag: source_eqtl_gtex_AdrenalGland
    provenance: GTEx
    version: v7
    url: https://www.gtexportal.org/home/datasets
    sample_origin: primary-tissue
    tissue: Adrenal Gland
    cell_type: mixed
    life_stage: adult
    sex: mixed
    species: human
  method:
    method_tag: soft_eqtl
    method_mode: software
    method_mode_ontology_term_id: ECO_0000203
    method_description: 'We cross-referenced the sentinel CAD variants and their proxies
      (r2≥0.8) with STARNET eQTLs

      reaching a 5% FDR for all tissues. To ensure the CAD association and eQTL are
      likely to be

      driven by the same causal variant, we retained only those eQTLs where the CAD-associated

      variant and the lead eQTL variant had r2≥0.8 among European ancestry participants
      from UK

      Biobank. '
- column_header: QTL_eQTL_gtex_Artery_Aorta_pvalue
  column_description: Nominal P-value for the association between genotype and gene
    expression in GTEx aortic artery tissue.
  evidence_stream_tag: eQTL
  evidence_category: Molecular QTL
  evidence_category_abbreviation: QTL
  variant_or_gene_centric: variant-centric
  source_tag: source_eqtl_gtex_ArteryAorta
  method_tag: soft_eqtl
  threshold: FDR<5%
  source:
    source_tag: source_eqtl_gtex_ArteryAorta
    provenance: GTEx
    version: v7
    url: https://www.gtexportal.org/home/datasets
    sample_origin: primary-tissue
    tissue: Artery Aorta
    cell_type: mixed
    life_stage: adult
    sex: mixed
    species: human
  method:
    method_tag: soft_eqtl
    method_mode: software
    method_mode_ontology_term_id: ECO_0000203
    method_description: 'We cross-referenced the sentinel CAD variants and their proxies
      (r2≥0.8) with STARNET eQTLs

      reaching a 5% FDR for all tissues. To ensure the CAD association and eQTL are
      likely to be

      driven by the same causal variant, we retained only those eQTLs where the CAD-associated

      variant and the lead eQTL variant had r2≥0.8 among European ancestry participants
      from UK

      Biobank. '
- column_header: QTL_eQTL_gtex_Artery_Coronary_pvalue
  column_description: Nominal P-value for the association between genotype and gene
    expression in GTEx coronary artery tissue.
  evidence_stream_tag: eQTL
  evidence_category: Molecular QTL
  evidence_category_abbreviation: QTL
  variant_or_gene_centric: variant-centric
  source_tag: source_eqtl_gtex_ArteryCoronary
  method_tag: soft_eqtl
  threshold: FDR<5%
  source:
    source_tag: source_eqtl_gtex_ArteryCoronary
    provenance: GTEx
    version: v7
    url: https://www.gtexportal.org/home/datasets
    sample_origin: primary-tissue
    tissue: Artery Coronary
    cell_type: mixed
    life_stage: adult
    sex: mixed
    species: human
  method:
    method_tag: soft_eqtl
    method_mode: software
    method_mode_ontology_term_id: ECO_0000203
    method_description: 'We cross-referenced the sentinel CAD variants and their proxies
      (r2≥0.8) with STARNET eQTLs

      reaching a 5% FDR for all tissues. To ensure the CAD association and eQTL are
      likely to be

      driven by the same causal variant, we retained only those eQTLs where the CAD-associated

      variant and the lead eQTL variant had r2≥0.8 among European ancestry participants
      from UK

      Biobank. '
- column_header: QTL_eQTL_gtex_Artery_Tibial_pvalue
  column_description: Nominal P-value for the association between genotype and gene
    expression in GTEx tibial artery tissue.
  evidence_stream_tag: eQTL
  evidence_category: Molecular QTL
  evidence_category_abbreviation: QTL
  variant_or_gene_centric: variant-centric
  source_tag: source_eqtl_gtex_ArteryTibial
  method_tag: soft_eqtl
  threshold: FDR<5%
  source:
    source_tag: source_eqtl_gtex_ArteryTibial
    provenance: GTEx
    version: v7
    url: https://www.gtexportal.org/home/datasets
    sample_origin: primary-tissue
    tissue: Artery Tibial
    cell_type: mixed
    life_stage: adult
    sex: mixed
    species: human
  method:
    method_tag: soft_eqtl
    method_mode: software
    method_mode_ontology_term_id: ECO_0000203
    method_description: 'We cross-referenced the sentinel CAD variants and their proxies
      (r2≥0.8) with STARNET eQTLs

      reaching a 5% FDR for all tissues. To ensure the CAD association and eQTL are
      likely to be

      driven by the same causal variant, we retained only those eQTLs where the CAD-associated

      variant and the lead eQTL variant had r2≥0.8 among European ancestry participants
      from UK

      Biobank. '
- column_header: QTL_eQTL_gtex_Liver_pvalue
  column_description: Nominal P-value for the association between genotype and gene
    expression in GTEx liver tissue.
  evidence_stream_tag: eQTL
  evidence_category: Molecular QTL
  evidence_category_abbreviation: QTL
  variant_or_gene_centric: variant-centric
  source_tag: source_eqtl_gtex_Liver
  method_tag: soft_eqtl
  threshold: FDR<5%
  source:
    source_tag: source_eqtl_gtex_Liver
    provenance: GTEx
    version: v7
    url: https://www.gtexportal.org/home/datasets
    sample_origin: primary-tissue
    tissue: Liver
    cell_type: mixed
    life_stage: adult
    sex: mixed
    species: human
  method:
    method_tag: soft_eqtl
    method_mode: software
    method_mode_ontology_term_id: ECO_0000203
    method_description: 'We cross-referenced the sentinel CAD variants and their proxies
      (r2≥0.8) with STARNET eQTLs

      reaching a 5% FDR for all tissues. To ensure the CAD association and eQTL are
      likely to be

      driven by the same causal variant, we retained only those eQTLs where the CAD-associated

      variant and the lead eQTL variant had r2≥0.8 among European ancestry participants
      from UK

      Biobank. '
- column_header: QTL_eQTL_gtex_Whole_Blood_pvalue
  column_description: Nominal P-value for the association between genotype and gene
    expression in GTEx whole blood.
  evidence_stream_tag: eQTL
  evidence_category: Molecular QTL
  evidence_category_abbreviation: QTL
  variant_or_gene_centric: variant-centric
  source_tag: source_eqtl_gtex_WholeBlood
  method_tag: soft_eqtl
  threshold: FDR<5%
  source:
    source_tag: source_eqtl_gtex_WholeBlood
    provenance: GTEx
    version: v7
    url: https://www.gtexportal.org/home/datasets
    sample_origin: primary-tissue
    tissue: Whole Blood
    cell_type: mixed
    life_stage: adult
    sex: mixed
    species: human
  method:
    method_tag: soft_eqtl
    method_mode: software
    method_mode_ontology_term_id: ECO_0000203
    method_description: 'We cross-referenced the sentinel CAD variants and their proxies
      (r2≥0.8) with STARNET eQTLs

      reaching a 5% FDR for all tissues. To ensure the CAD association and eQTL are
      likely to be

      driven by the same causal variant, we retained only those eQTLs where the CAD-associated

      variant and the lead eQTL variant had r2≥0.8 among European ancestry participants
      from UK

      Biobank. '
- column_header: QTL_eQTL_STARNET_AOR_adjusted_pvalue
  column_description: Adjusted P-value for the association between genotype and gene
    expression in atherosclerotic aortic root tissue from STARNET.
  evidence_stream_tag: eQTL
  evidence_category: Molecular QTL
  evidence_category_abbreviation: QTL
  variant_or_gene_centric: variant-centric
  source_tag: source_eqtl_starnet_AOR
  method_tag: soft_eqtl
  threshold: FDR<5%
  source:
    source_tag: source_eqtl_starnet_AOR
    provenance: STARNET
    doi: 10.1126/science.aad6970
    sample_origin: primary-tissue
    tissue: Atherosclerotic Aortic Root
    cell_type: mixed
    disease: disease
    life_stage: adult
    sex: mixed
    species: human
    description: STARNET tissue from donors with coronary artery disease.
  method:
    method_tag: soft_eqtl
    method_mode: software
    method_mode_ontology_term_id: ECO_0000203
    method_description: 'We cross-referenced the sentinel CAD variants and their proxies
      (r2≥0.8) with STARNET eQTLs

      reaching a 5% FDR for all tissues. To ensure the CAD association and eQTL are
      likely to be

      driven by the same causal variant, we retained only those eQTLs where the CAD-associated

      variant and the lead eQTL variant had r2≥0.8 among European ancestry participants
      from UK

      Biobank. '
- column_header: QTL_eQTL_STARNET_BLOOD_adjusted_pvalue
  column_description: Adjusted P-value for the association between genotype and gene
    expression in blood from STARNET.
  evidence_stream_tag: eQTL
  evidence_category: Molecular QTL
  evidence_category_abbreviation: QTL
  variant_or_gene_centric: variant-centric
  source_tag: source_eqtl_starnet_BLOOD
  method_tag: soft_eqtl
  threshold: FDR<5%
  source:
    source_tag: source_eqtl_starnet_BLOOD
    provenance: STARNET
    doi: 10.1126/science.aad6970
    sample_origin: primary-tissue
    tissue: Blood
    cell_type: mixed
    disease: disease
    life_stage: adult
    sex: mixed
    species: human
    description: STARNET tissue from donors with coronary artery disease.
  method:
    method_tag: soft_eqtl
    method_mode: software
    method_mode_ontology_term_id: ECO_0000203
    method_description: 'We cross-referenced the sentinel CAD variants and their proxies
      (r2≥0.8) with STARNET eQTLs

      reaching a 5% FDR for all tissues. To ensure the CAD association and eQTL are
      likely to be

      driven by the same causal variant, we retained only those eQTLs where the CAD-associated

      variant and the lead eQTL variant had r2≥0.8 among European ancestry participants
      from UK

      Biobank. '
- column_header: QTL_eQTL_STARNET_LIV_adjusted_pvalue
  column_description: Adjusted P-value for the association between genotype and gene
    expression in liver tissue from STARNET.
  evidence_stream_tag: eQTL
  evidence_category: Molecular QTL
  evidence_category_abbreviation: QTL
  variant_or_gene_centric: variant-centric
  source_tag: source_eqtl_starnet_LIV
  method_tag: soft_eqtl
  threshold: FDR<5%
  source:
    source_tag: source_eqtl_starnet_LIV
    provenance: STARNET
    doi: 10.1126/science.aad6970
    sample_origin: primary-tissue
    tissue: Liver
    cell_type: mixed
    disease: disease
    life_stage: adult
    sex: mixed
    species: human
    description: STARNET tissue from donors with coronary artery disease.
  method:
    method_tag: soft_eqtl
    method_mode: software
    method_mode_ontology_term_id: ECO_0000203
    method_description: 'We cross-referenced the sentinel CAD variants and their proxies
      (r2≥0.8) with STARNET eQTLs

      reaching a 5% FDR for all tissues. To ensure the CAD association and eQTL are
      likely to be

      driven by the same causal variant, we retained only those eQTLs where the CAD-associated

      variant and the lead eQTL variant had r2≥0.8 among European ancestry participants
      from UK

      Biobank. '
- column_header: QTL_eQTL_STARNET_MAM_adjusted_pvalue
  column_description: Adjusted P-value for the association between genotype and gene
    expression in internal mammary artery tissue from STARNET.
  evidence_stream_tag: eQTL
  evidence_category: Molecular QTL
  evidence_category_abbreviation: QTL
  variant_or_gene_centric: variant-centric
  source_tag: source_eqtl_starnet_MAM
  method_tag: soft_eqtl
  threshold: FDR<5%
  source:
    source_tag: source_eqtl_starnet_MAM
    provenance: STARNET
    doi: 10.1126/science.aad6970
    sample_origin: primary-tissue
    tissue: Atherosclerotic-lesion-free Internal Mammary Artery
    cell_type: mixed
    disease: disease
    life_stage: adult
    sex: mixed
    species: human
    description: STARNET tissue from donors with coronary artery disease.
  method:
    method_tag: soft_eqtl
    method_mode: software
    method_mode_ontology_term_id: ECO_0000203
    method_description: 'We cross-referenced the sentinel CAD variants and their proxies
      (r2≥0.8) with STARNET eQTLs

      reaching a 5% FDR for all tissues. To ensure the CAD association and eQTL are
      likely to be

      driven by the same causal variant, we retained only those eQTLs where the CAD-associated

      variant and the lead eQTL variant had r2≥0.8 among European ancestry participants
      from UK

      Biobank. '
- column_header: QTL_eQTL_STARNET_SF_adjusted_pvalue
  column_description: Adjusted P-value for the association between genotype and gene
    expression in subcutaneous fat tissue from STARNET.
  evidence_stream_tag: eQTL
  evidence_category: Molecular QTL
  evidence_category_abbreviation: QTL
  variant_or_gene_centric: variant-centric
  source_tag: source_eqtl_starnet_SF
  method_tag: soft_eqtl
  threshold: FDR<5%
  source:
    source_tag: source_eqtl_starnet_SF
    provenance: STARNET
    doi: 10.1126/science.aad6970
    sample_origin: primary-tissue
    tissue: Subcutaneous Fat
    cell_type: mixed
    disease: disease
    life_stage: adult
    sex: mixed
    species: human
    description: STARNET tissue from donors with coronary artery disease.
  method:
    method_tag: soft_eqtl
    method_mode: software
    method_mode_ontology_term_id: ECO_0000203
    method_description: 'We cross-referenced the sentinel CAD variants and their proxies
      (r2≥0.8) with STARNET eQTLs

      reaching a 5% FDR for all tissues. To ensure the CAD association and eQTL are
      likely to be

      driven by the same causal variant, we retained only those eQTLs where the CAD-associated

      variant and the lead eQTL variant had r2≥0.8 among European ancestry participants
      from UK

      Biobank. '
- column_header: QTL_eQTL_STARNET_SKLM_adjusted_pvalue
  column_description: Adjusted P-value for the association between genotype and gene
    expression in skeletal muscle tissue from STARNET.
  evidence_stream_tag: eQTL
  evidence_category: Molecular QTL
  evidence_category_abbreviation: QTL
  variant_or_gene_centric: variant-centric
  source_tag: source_eqtl_starnet_SKLM
  method_tag: soft_eqtl
  threshold: FDR<5%
  source:
    source_tag: source_eqtl_starnet_SKLM
    provenance: STARNET
    doi: 10.1126/science.aad6970
    sample_origin: primary-tissue
    tissue: Skeletal Muscle
    cell_type: mixed
    disease: disease
    life_stage: adult
    sex: mixed
    species: human
    description: STARNET tissue from donors with coronary artery disease.
  method:
    method_tag: soft_eqtl
    method_mode: software
    method_mode_ontology_term_id: ECO_0000203
    method_description: 'We cross-referenced the sentinel CAD variants and their proxies
      (r2≥0.8) with STARNET eQTLs

      reaching a 5% FDR for all tissues. To ensure the CAD association and eQTL are
      likely to be

      driven by the same causal variant, we retained only those eQTLs where the CAD-associated

      variant and the lead eQTL variant had r2≥0.8 among European ancestry participants
      from UK

      Biobank. '
- column_header: QTL_eQTL_STARNET_VAF_adjusted_pvalue
  column_description: Adjusted P-value for the association between genotype and gene
    expression in visceral abdominal fat tissue from STARNET.
  evidence_stream_tag: eQTL
  evidence_category: Molecular QTL
  evidence_category_abbreviation: QTL
  variant_or_gene_centric: variant-centric
  source_tag: source_eqtl_starnet_VAF
  method_tag: soft_eqtl
  threshold: FDR<5%
  source:
    source_tag: source_eqtl_starnet_VAF
    provenance: STARNET
    doi: 10.1126/science.aad6970
    sample_origin: primary-tissue
    tissue: Visceral Abdominal Fat
    cell_type: mixed
    disease: disease
    life_stage: adult
    sex: mixed
    species: human
    description: STARNET tissue from donors with coronary artery disease.
  method:
    method_tag: soft_eqtl
    method_mode: software
    method_mode_ontology_term_id: ECO_0000203
    method_description: 'We cross-referenced the sentinel CAD variants and their proxies
      (r2≥0.8) with STARNET eQTLs

      reaching a 5% FDR for all tissues. To ensure the CAD association and eQTL are
      likely to be

      driven by the same causal variant, we retained only those eQTLs where the CAD-associated

      variant and the lead eQTL variant had r2≥0.8 among European ancestry participants
      from UK

      Biobank. '
- column_header: DB_ClinVar
  column_description: 'List of genes in CAD-associated regions for which pathogenic
    (or likely pathogenic) variants were found in ClinVar for phenotypes and disorders
    related to CAD. '
  evidence_category: Association from curated database
  evidence_category_abbreviation: DB
  variant_or_gene_centric: gene-centric
  source_tag: source_clinvar
  method_tag: method_clinvar
  note: The report disease to the gene in the region harbors a variant that ClinVar
    classifies as having evidence for being pathogenic for a cardiovascular-relevant
    monogenic disorder
  source:
    source_tag: source_clinvar
    provenance: 'ClinVar '
    version: 2020-06-06 00:00:00
    url: https://www.ncbi.nlm.nih.gov/clinvar/
  method:
    method_tag: method_clinvar
    method_mode: manual
    method_mode_ontology_term_id: ECO_0000218
    method_description: Genes associated with cardiovascular-related pathogenic or
      likely pathogenic variants in ClinVar within ±500 kb of CAD sentinel variants.
      Variants linked to multiple genes were assigned to all associated genes.
- column_header: LIT_RareVariant
  column_description: 'A gene in the region has been shown to have a rare variant
    association with CAD in a previous WES or genotyping study '
  evidence_category: Literature curation
  evidence_category_abbreviation: LIT
  variant_or_gene_centric: gene-centric
  note: "reviewed the literature for three \nsources of evidence as follows: rare\
    \ coding variants previously \nassociated with CAD, either individually or in\
    \ aggregate gene-based \ntests, through whole-exome sequencing (WES) or exome\
    \ array stud-\nies;"
- column_header: LIT_MRorDrug
  column_description: A gene in the region has been implicated by an effective drug
    targeting the protein and/or a positive MR study suggesting a causal effect of
    the protein on CAD
  evidence_category: Literature curation
  evidence_category_abbreviation: LIT
  variant_or_gene_centric: gene-centric
  note: "reviewed the literature for three \nsources of evidence as follows: Mendelian\
    \ randomization (MR) studies of gene expression, \nprotein levels or proximal\
    \ phenotypes that implicate specific genes as \ncausal effector genes for CAD\
    \ and drugs proven to be effective for \ncardiovascular-relevant indications and\
    \ that target specific proteins \nencoded by genes."
- column_header: PERTURB_mouse_modelId
  column_description: Specifies the mouse model or experimental system used to test
    gene perturbation effects, particularly those leading to cardiovascular-relevant
    phenotypes.
  evidence_category: Perturbation
  evidence_category_abbreviation: PERTURB
  variant_or_gene_centric: gene-centric
  source_tag: source_perturb_mouse
  note: List of cardiovascular-relevant mouse phenotypes for gene knockouts of genes
    in genome-wide significant association regions. Columns describe the gene (including
    HGNC and Mouse Genome Informatics IDs), the mouse model ID and description, and
    the mouse phenotyping ID and description. The table is ordered alphabetically
    by gene name.
  source:
    source_tag: source_perturb_mouse
    provenance: Mouse Genome Informatics database
    version: 2019
    url: http://www.informatics.jax.org
- column_header: PERTURB_mouse_phenotype
  column_description: a gene for which a mouse knock-out has a cardiovascular-relevant
    phenotype
  evidence_category: Perturbation
  evidence_category_abbreviation: PERTURB
  variant_or_gene_centric: gene-centric
  source_tag: source_perturb_mouse
  note: List of cardiovascular-relevant mouse phenotypes for gene knockouts of genes
    in genome-wide significant association regions. Columns describe the gene (including
    HGNC and Mouse Genome Informatics IDs), the mouse model ID and description, and
    the mouse phenotyping ID and description. The table is ordered alphabetically
    by gene name.
  source:
    source_tag: source_perturb_mouse
    provenance: Mouse Genome Informatics database
    version: 2019
    url: http://www.informatics.jax.org
Integration:
- integration_tag: pops_500
  column_header: INT_PoPS_500kb_score
  column_description: PoPS score for all genes within a 500kb window of each of the
    279 sentinel genome-wide significant associations with CAD
  author_conclusion: false
  method_tag: soft_pops
  note: ' gene with highest PoPS score within 500kb of sentinel.              '
  method:
    method_tag: soft_pops
    method_mode: software
    method_mode_ontology_term_id: ECO_0007674
    software_name: PoPS
    software_doi: https://doi.org/10.1101/2020.09.08.20190561
- integration_tag: pops_500
  column_header: INT_PoPS_500kb_rank
  column_description: PoPS association-specific rank for all genes within a 500kb
    window of each of the 279 sentinel genome-wide significant associations with CAD
  author_conclusion: false
  method_tag: soft_pops
  method:
    method_tag: soft_pops
    method_mode: software
    method_mode_ontology_term_id: ECO_0007674
    software_name: PoPS
    software_doi: https://doi.org/10.1101/2020.09.08.20190561
- integration_tag: pops_500
  column_header: INT_PoPS_500kb_summary
  column_description: Binary indicator of PoPS support. Equals 1 if the gene is ranked
    among the top 2 PoPS-prioritized genes within ±500 kb of the locus, providing
    positive evidence that the gene may be causal; otherwise 0.
  author_conclusion: false
  method_tag: soft_pops
  method:
    method_tag: soft_pops
    method_mode: software
    method_mode_ontology_term_id: ECO_0007674
    software_name: PoPS
    software_doi: https://doi.org/10.1101/2020.09.08.20190561
- integration_tag: n_predictors
  column_header: INT_n_predictors
  column_description: Number of evidence streams supporting the variant-gene pair
    for the phenotype.
  author_conclusion: false
  evidence_streams_included: PROX | QTL | FUNC | LIT | PERTURB | DB
  integrations_included: pops
  method_tag: method_customised
  note: Unweighted count of the eight evidence predictors described by the publication.
  method:
    method_tag: method_customised
    method_mode: manual
    method_mode_ontology_term_id: ECO_0007674
    method_description: 'To systematically prioritize likely causal genes for all
      279 genome-wide associations, we integrated the following eight of the aforementioned
      similarity-based or locus-based predictors of causal genes: (1) the top two
      prioritized genes from PoPS; (2) genes with eQTLs in CAD-relevant tissues from
      STARNET or GTEx; (3) genes containing protein-altering variants that are in
      strong LD (r2 ≥ 0.8) with the CAD sentinel variant; (4) genes harboring variants
      responsible for monogenic disorders of cardiovascular relevance according to
      ClinVar; (5) genes containing rare coding variants that have been associated
      with CAD risk in previous WES or array-based studies; (6) genes encoding proteins
      of causal relevance to CAD per MR studies or that are targets for established
      cardiovascular drugs; (7) genes that display cardiovascular-relevant phenotypes
      in knock-out mice from the International Mouse Phenotyping Consortium or Mouse
      Genome Informatics database; and (8) the nearest gene to the CAD sentinel variant
      (Fig. 5a). We prioritized the most likely ‘causal gene’ for each association
      using a consensus-based approach, selecting the gene with the highest, unweighted
      sum of evidence across all eight predictors.'
- integration_tag: author_conclusion
  column_header: INT_author_conclusion
  column_description: Whether this variant-gene pair appears in the 44-gene display
    subset in Aragam et al. Figure 5b.
  author_conclusion: true
  integrations_included: n_predictors
  method_tag: method_customised
  threshold: 'Displayed in Figure 5b: at least four agreeing predictors; one sentinel
    association shown per gene.'
  method:
    method_tag: method_customised
    method_mode: manual
    method_mode_ontology_term_id: ECO_0007674
    method_description: 'To systematically prioritize likely causal genes for all
      279 genome-wide associations, we integrated the following eight of the aforementioned
      similarity-based or locus-based predictors of causal genes: (1) the top two
      prioritized genes from PoPS; (2) genes with eQTLs in CAD-relevant tissues from
      STARNET or GTEx; (3) genes containing protein-altering variants that are in
      strong LD (r2 ≥ 0.8) with the CAD sentinel variant; (4) genes harboring variants
      responsible for monogenic disorders of cardiovascular relevance according to
      ClinVar; (5) genes containing rare coding variants that have been associated
      with CAD risk in previous WES or array-based studies; (6) genes encoding proteins
      of causal relevance to CAD per MR studies or that are targets for established
      cardiovascular drugs; (7) genes that display cardiovascular-relevant phenotypes
      in knock-out mice from the International Mouse Phenotyping Consortium or Mouse
      Genome Informatics database; and (8) the nearest gene to the CAD sentinel variant
      (Fig. 5a). We prioritized the most likely ‘causal gene’ for each association
      using a consensus-based approach, selecting the gene with the highest, unweighted
      sum of evidence across all eight predictors.'
```
