## Supplemental File 3 for "Predicted Effector Gene Aggregation, Standards and Unified Schema (PEGASUS): A Community Framework for Effector Gene Reporting": PEG_dummy_submission_data_explanation.docx

### PEGASUS Dummy Submission data description

*As a dummy submission, please note that it is* ***NOT*** *a full dataset. It only includes locus 6 here for testing purposes.*

1. What files are included in this dummy submission:
   a. matrix_Aragam_PEGSt000007.tsv

- Contains the PEG matrix, where each row represents a unique variant–gene pair and includes evidence values associated with the variant, the gene, or the variant→gene relationship.
- This file provides the full raw dataset, allowing downstream data analysts to reuse or reprocess the complete evidence matrix.
  b. list_Aragam_PEGSt000007.tsv
- Contains the prioritised gene list together with the availability of different evidence types.
- Most columns are represented as True/False, indicating whether a given evidence type is present.
- Some fields — particularly the conclusion column — may contain free-text or other data types.
  c. metadata_Aragam_PEGSt000007.xlsx
- An Excel-based metadata template designed to be easy for data submitters to complete.
- It captures essential study information, contributor details, data provenance, and other required fields.
  d. metadata_Aragam_PEGSt000007.yml
- A machine-readable metadata file, generated by converting the completed metadata template into YAML format.
- This file is intended for data users and backend systems to understand the metadata context.
- Although YAML is used here for demonstration, the metadata could be stored in other formats depending on system requirements.

2. What may happen during submission
   a. What the user provides
      i. PEG matrix file
      ii. metadata template
   b. What the backend performs
      i. Data validation — checks format, required fields, and identifiers. (https://ebispot.github.io/PEGASUS/docs/peg-matrix/)
      ii. Metadata validation — verifies that all mandatory fields in the template are correctly filled and data type.

(https://ebispot.github.io/PEGASUS/docs/peg-metadata/)

iii. Metadata conversion — transforms the validated metadata template into a structured YAML file (or any other format) for ingestion. (Yue can help with it to prepare a Python script)
iv. Extract the PEG List from the PEG matrix. The top gene in each locus.

3. How the data is presented on the UI:
   a. The final presentation of PEG matrix data, prioritised PEG gene lists, and study metadata depends on the design goals of the portal.
